# The effects of electroencephalogram feature-based transcranial alternating current stimulation on working memory and electrophysiology

**DOI:** 10.1101/2022.01.19.476885

**Authors:** Lanting Zeng, Mingrou Guo, Ruoling Wu, Yu Luo, Pengfei Wei

## Abstract

Transcranial alternating current stimulation (tACS) can influence cognitive functions by modulating brain oscillations. However, results regarding the effectiveness of tACS in regulating cognitive performance have been inconsistent. In the present study, we aimed to find EEG characteristics associated with the improvements in working memory performance, to select tACS stimulus targets and frequency based on this feature, and to explore effects of selected stimulus on verbal working memory. To achieve this goal, we first investigated the EEG characteristics associated with improvements in working memory performance with the aid of EEG analyses and machine learning techniques. These analyses suggested that 8 Hz activity in the prefrontal region was related to accuracy in the verbal working memory task. The tACS stimulus target and pattern were then selected based on the EEG feature. Finally, the selected tACS frequency (8 Hz tACS in the prefrontal region) was applied to modulate working memory. The performance of working memory was improved significantly using the selected stimulation than using 40 Hz and sham stimulation (Especially for participants with low verbal working memory). In conclusion, using EEG features related to positive behavioral changes to select brain regions and stimulation patterns for tACS is an effective intervention for improving working memory. Our results contribute to the groundwork for future tACS closed-loop interventions for cognitive deterioration.

## 1 Introduction

Over the past few decades, the development of non-invasive brain stimulation (NIBS) techniques has provided a new and effective approach to modulate memory for both researchers and clinicians (Misselhorn et al., 2020; Reinhart & Nguyen, 2019; Rombouts et al., 2005; Benussi et al., 2021; Grover et al., 2021). Among NIBS techniques, transcranial alternating current stimulation (tACS) can alter specific frequencies of brain oscillations in predefined brain regions and further modulate human cognition (Zaehle et al., 2010; Vosskuhl et al., 2015; Riddle et al., 2021). Working memory deterioration is a key feature of cognitive decline in old age (Li et al., 2001). Although some researchers have proposed that NIBS can help to regulate memory and attenuate age-related cognitive decline (Reinhart & Nguyen, 2019), results regarding the effectiveness of tACS in regulating working memory performance have been inconsistent. Given that verbal and visual working memory involve different cognitive structures, these inconsistencies may have been due to improper selection of stimulation targets and parameters. Therefore, in the current study, we want to select tACS stimulus targets and frequency based on the EEG characteristics associated with improvements in verbal working memory, and to explore the effect of selected stimulus on verbal working memory.

Several studies have indicated that theta and gamma tACS can improve verbal working memory. Based on the positive association between gamma band activity and task performance reported in previous studies, Hoy et al. (2015) applied 40 Hz tACS to the F3-contralateral supraorbital area in 18 healthy participants. Participants underwent 20 min of tES (40 Hz or sham) while completing a verbal two-back task, as well as two-back and three-back tasks before and after tACS. Compared with sham-tACS and transcranial direct current stimulation (tDCS), 40 Hz tACS resulted in increased performance in terms of d prime (an accuracy discriminability index). Biel et al. (2021) also recently reported that frontoparietal in-phase and in-phase focal theta tACS substantially improved verbal three-back task performance when compared with placebo stimulation.

However, some studies have reported that tACS was not effective or was only effective in a limited number of people for verbal working memory. For example, Pahor & Jaušovec (2018) applied tACS over many regions (F3-F4, F3-P3, F4-P4, P3-P4) in healthy adults to investigate working memory using two-back and three-back tasks. The rationale of the electrode montage and frequency band was based on previous correlational research, which showed that frontotemporal theta and gamma frequency bands are involved in working memory. Nevertheless, only theta-tACS improved performance on the three-back task in the F4-P4 region. In an earlier study, Vosskuhl et al. (2015) applied individual theta frequency stimulation at Pz-FPz. When compared with sham stimulation, tACS was associated with improved short-term memory performance. However, there was no significant difference in improvements on the verbal three-back task between tACS and sham stimulation. Kilian et al. (2020) further compared the effects of tDCS and 6-Hz tACS applied at F3-FP2 in healthy participants, reporting no significant difference in verbal n-back task performance among the experimental groups (sham, tDCS, and tACS), but they observed that tDCS and tACS exert different modulatory effects on fMRI-derived network dynamics.

In the abovementioned studies, stimulation targeted the prefrontal, frontal, and parietal lobes using theta and gamma frequencies. In NIBS studies, specific targets and parameters for stimulation are usually selected in the following two ways: (a) frequency bands and regions are determined based on previously reported findings regarding their association with verbal working memory or (b) the parameters are simply selected based on those used in previous studies. While these methods have been somewhat successful, there is no guarantee that each combination of parameters will regulate working memory.

We hypothesized that after identifying the brain regions and frequency bands associated with working memory, further exploration of changes in electroencephalogram (EEG) activity that correspond to positive behavioral changes can help to improve the effectiveness of tACS by enabling researchers to set stimulation targets and parameters based on such EEG activity. Repeated assessments of verbal working memory and EEG activity may therefore help to elucidate the electrophysiological features that vary with improvements in behavioral performance. To test this hypothesis, we conducted two experiments that mainly focused on working memory. Experiment 1 was an EEG study, wherein participants completed three n-back tasks, and the electrophysiological features related to improvements in working memory were extracted. In Experiment 2, the participants were divided into three groups and received different frequencies of online tACS: the frequency in group 1 was the evident band in experiment 1, the frequency in group 2 was the non-evident band in experiment 1, and group 3 was the sham group. The results of the comparison between group 1 and the other groups can answer the research question.

## 2 Experiment 1: EEG features related to performance

### 2.1 Materials and Methods

#### 2.1.1 Participants

A total of 35 healthy adults aged 22–26 years of age participated in Experiment 1. All participants had normal or corrected-to-normal vision and were right-handed.

In experiment 1, ten participants were excluded because they did not complete the experiment and 25 participants (five females; mean age 23.76±1.14 years) were included in the analyses.

When analyzing the results of Experiment 1, we considered that some volunteers would exhibit naturally high performance on the verbal working memory task, leading to a ceiling effect over multiple measurements that may impede identification of the EEG characteristics associated with improvements in performance. We also considered that individuals with high and low levels of verbal working memory ability may exhibit differences in EEG activity and that the same tES parameter may exert different modulatory effects in each group (Daffner et al., 2011, Tseng et al., 2012). For the behavioral analyses, participants were divided into two groups based on their performance in block 1. The grouping method was selected in reference to previous studies (Daffner et al., 2011, Tseng et al., 2012). The scores of the three-back and four-back tasks were summed. The participants who scored lower than the median scores were assigned to the low-performance group, while those who scored higher than the median scores were assigned to the high-performance group. Following grouping, four participants were excluded due to extreme values (target accuracy [target-ACC] or reaction time (RT) exceeding two standard deviations from the mean). The final low-performance group (LP) and high-performance groups (HP) included nine and 12 participants, respectively.

For the EEG analyses, two participants were excluded because they had not sufficient number of good quality EEG trials after artifact removal (LP group: n = 8; HP group: n = 11).

This study was approved by the Ethics Committee of the Shenzhen Institute of Advanced Technology. The experimental procedures conformed to the principles of the Declaration of Helsinki regarding human experimentation. All participants provided oral consent, signed informed consent documents, and received 270RMB for their participation.

#### 2.1.2 Experimental Design and Schedule

In Experiment 1, all subjects received the same treatment. Participants were required to visit the laboratory twice to complete three n-Back tasks (blocks 1, 2, and 3). On day 1, participants performed the block 1 n-back task. After 1 week, the subjects returned to the laboratory and completed blocks 2 and 3. There was a 10 min break between blocks 2 and 3. In each task, task-state EEG data were recorded.

#### 2.1.3 N-back Task

Kirchner (1958) first proposed the n-back task. Subsequently, the n-back task has been widely employed to investigate and measure working memory. In Experiment 1, we employed the two-back task as an exercise, and the three-back and four-back tasks to measure the working memory performance of volunteers. As illustrated in Figure 1, in n-Back task (e.g., two-back task), each trial started with a stimulus consisting of an uppercase letter presented for 2 s, followed by a fixation “+” for 0.5 s. After the n^th^ trial (e.g., 2^nd^ in the two-back task), participants were required to determine whether the current letter was the same as the previous n^th^ letter (e.g., 2^nd^ in the two-back task). If they were the same, the participants were required to press the ‘match’ button. The current trial was defined as a target trial. Otherwise, the participants pressed the ‘non-match’ button, and the current trial was defined as a non-target trial. The accuracy of the target trials is defined as “target-ACC”. For each trial, participants had 2.5 s to respond and were instructed to press the button as quickly as possible. The instructions were similar in the three-back and four-back tasks.

**Figure 1.**
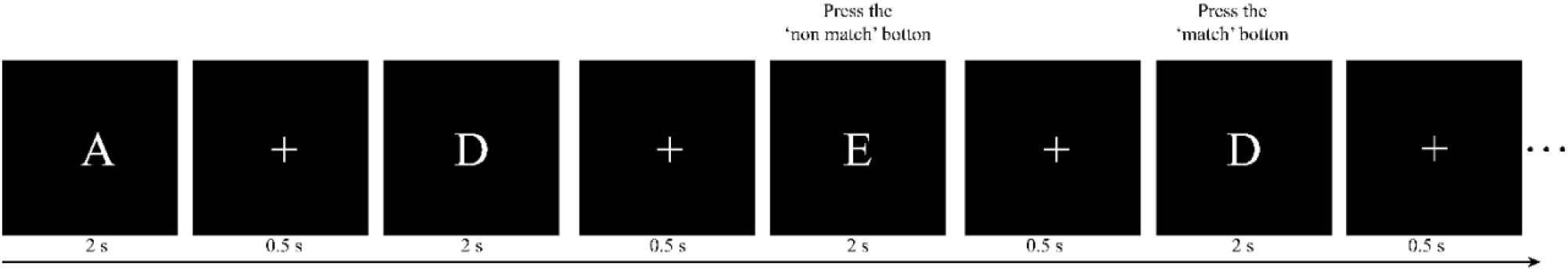
Illustration of the two-back task paradigm in this study. For the first two letters, participants were not required to press buttons, but keep the letters in their mind instead. Subsequently, for each letter, participants were required to determine whether the current letter was the same as the previous. In this case, the third letter should be compared with the first (“E” vs. “A”: non-match) and the fourth should be compared with the second one (“D” vs. “D”: match).

Each load condition (three-back and four-back) had one sequence of 60+n trials. Each sequence consisted of 20 trials for targets and 40 trials for non-targets. To help participants understand the n-back task requirements, practice trials were provided for each task. Each n-back task took 10-15 minutes to complete. The paradigms were programmed in MATLAB using PsychToolbox (Brainard, 1997; Pelli, 1997).

#### 2.1.4 Electrophysiological Recordings

The EEG was recorded during each n-back task with an online reference against the CPz electrode using a 64-channel wireless EEG amplifier with a sampling rate of 1000 Hz (NeuSen. W64, Neuracle, Changzhou, China). The ground electrode was located on the forehead (between the FPz and Fz electrodes). Electrode impedances were maintained at <5 kΩ.

#### 2.1.5 Initial EEG analysis

Initial EEG analysis includes two steps: (a) EEG signal preprocessing to remove artifacts and to improve the reliability of data and; (b) Preliminary exploration of brain regions and frequency bands with the activity corresponding to improvements in performance.

For EEG signal preprocessing, all data were analyzed using EEGLAB version 13.0.0b running in MATLAB (The MathWorks, USA). Only correctly responded trials were used in the analysis. Preprocessing steps included filtering (1-48 Hz), epoching (1000 ms before and 1500 ms after stimulus onset), baseline correction (500 ms before stimulus onset), and large artifact removal. Ocular artifacts were removed from the independent component analysis (ICA) results. The EEG data were then average-referenced. Finally, epochs that contained signals >100 μV from baseline were rejected.

In the second step, we used the function *pop_newtimef* (Arnaud Delorme, CNL/Salk Institute, 2001) in EEGLAB for time-frequency analysis. To compare the changes of EEG activity between block 1 and block 3. The number of cycles in each analysis wavelet was [3 0.5], the padratio was 2, and the window length was 350 ms. Meanwhile, the filter bank common spatial pattern (FBCSP) was used to explore spatio-frequency modes corresponding to improvements in performance.

FBCSP is a machine learning approach used to extract the optimal spatial features from different frequency bands (Ang et al., 2008). The original FBCSP algorithm consists of four steps: (1) band filtering, (2) spatial filtering, (3) mutual information (MI)-based feature selection, and (4) classification. MI is a useful statistical measure that can be used to quantify the relationship between variables (Timme & Lapish, 2018). Here, we dropped the classification step. Instead, we focused on the spatio-frequency modes (i.e., the brain regions and frequency bands) of the selected features. Figure 2 illustrates the workflow. To begin this process, FIR band-pass filters were employed to filter the EEG signals into three frequency bands: theta (4-8 Hz), alpha (8-13 Hz), and beta (13-30 Hz). 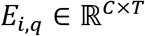 denotes the i-th trial of the qth frequency band EEG. In the spatial filtering step, we first calculated a spatial filter *W_i,q_* for each frequency band using the CSP algorithm (Blankertz et al., 2007; Pfurtscheller & Neuper, 2001). Notably, 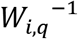 is the spatial distribution pattern of EEG signals. The spatial filter *W_i,q_* was then applied to the EEG matrix *E_i,q_*,

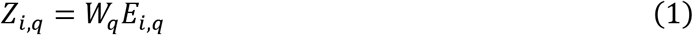

where the projected EEG matrix is 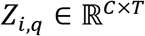. We selected the m first and rows of *Z_i,q_* to maximize the variation for one class while minimizing the variance for the other class. The normalized feature vector 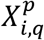 was then computed as follows:

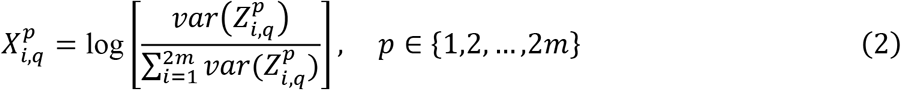

**Figure 2.**
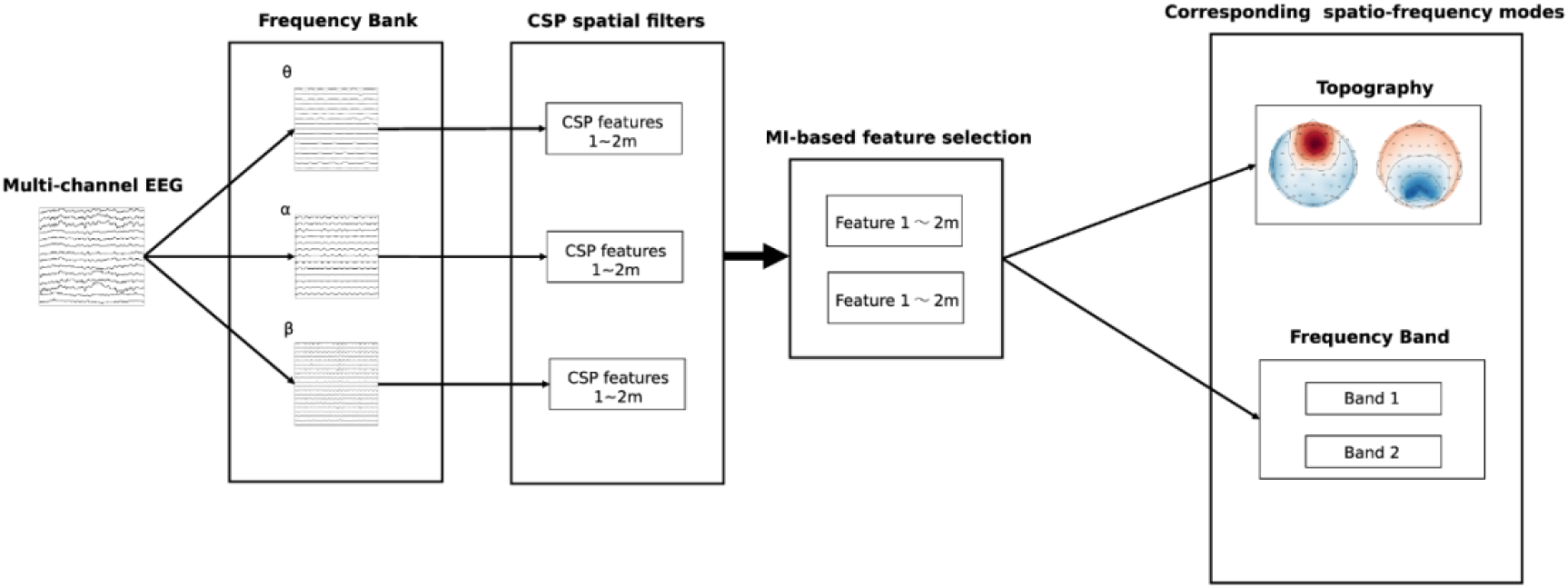
The workflow of filter bank common spatial pattern-based spatio-frequency mode selection. We first filtered the raw electroencephalogram into three frequency bands and then performed spatial filtering to obtain the common spatial pattern (CSP) features. Based on the mutual information, we selected the two most discriminate features and determined their associated spatio-frequency modes.

In the third step, the MI-based feature selection method was adopted to find the spatio-frequency modes containing the most discriminating features (Battiti, 1994). We defined the binary labels set as *l* ∈ *L* = {0,1}, where label 0 is for the lower-capacity subjects and label 1 is for the higher-capacity subjects. The mutual information 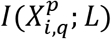 (MI-value) was defined as (Cover, 1999):

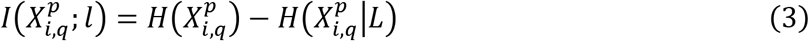

where the entropy for the T-dimensional feature vector 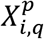 is

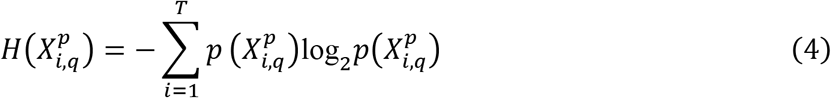

and the conditional entropy for the random variable 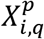 and *L* is

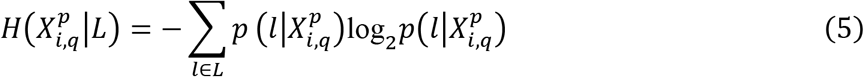

We selected the top two largest MI values for each n-back test. The corresponding brain regions and frequency bands were considered the most important spatio-frequency modes for n-back performance discrimination.

#### 2.1.6 Graph Convolutional Neural Network (GCNN)

We adapted the original Graph Convolutional Neural Network (GCNN) by adding an attention layer to capture brain network dynamics and identify the channel providing the greatest contribution to the n-back tasks. The GCNN is a generalized version of the convolutional neural network (CNN) (Defferrard et al., 2016). By employing spectral graph theory (Chung & Graham, 1997), GCNN can reveal the underlying topological information of high-dimensional data. In the second step, we investigated the intrinsic spatial patterns of multichannel EEG data using a GCNN model, in which each vertex represents an EEG channel and each edge represents the connection between two electrodes. Although the GCNN approach is effective for elucidating the spatial patterns of multichannel EEG, one limitation is the requirement for a fixed graph representation. In other words, the adjacent matrix must be predetermined before applying the GCNN to the data. However, the brain states of participants can exhibit time variance during long recording periods. Consequently, inspired by graph attention network (GAT) methods (Veličković et al., 2017), we adapted the original GCNN by adding an attention layer to capture brain network dynamics and identify the channel providing the greatest contribution in the n-back tasks.

By definition, a graph can be represented as *G* = {*V,E,A*}, in which *V* is the set of vertices with the number of *N* = |*V*|. *A* represents the adjacent matrix, in which each entry denotes the connection relationship (i.e., the edge) between two vertices. The set of input features can be denoted as 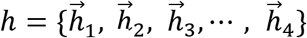, where each feature vector corresponds to a vertex. We first initialized the adjacent matrix randomly. The initial adjacent matrix can be updated by the graph attention layer (Veličković et al., 2017) during the training process. The updating rule is presented as follows:

First, the graph attention layer computes the attention coefficient matrix 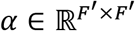, where *F′* is the size of the output feature set. The coefficients can be computed as

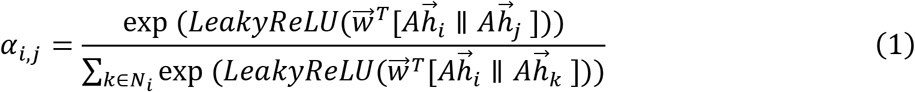

where *N_i_* is the set of adjacent vertices of the vertex *i*, 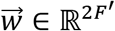 is the parameter vector of the graph attention layer, and ║ is the concatenation operation.

The adjacent matrix *A* can be updated by multiplying the coefficient matrix and the original adjacent matrix, as follows:

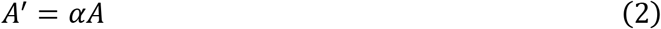

Meanwhile, the graph attention layer also updates the feature set according to the following:

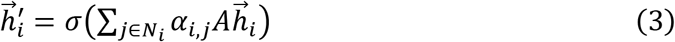

Then, two GCNN layers are used to classify the performance of the participants. *L* denotes the Laplacian matrix, which can be written as

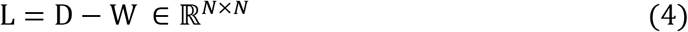

where *D* represents the degree matrix. *L* can then be decomposed as follows:

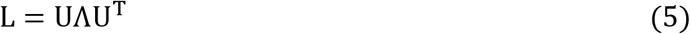

The convolution in the non-Euclidean domains can be computed as

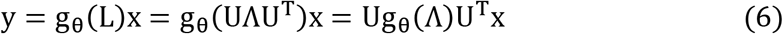

where g_θ_ is the non-parametric filter with learnable parameters. A fully connected layer is then adopted to predict behavioral performance.

#### 2.1.7 Further EEG Analysis

Further EEG analysis based on the results of initial EEG analysis and GCNN, which we explored the frequency change (4 Hz, 5 Hz, 6 Hz, 7 Hz, 8 Hz) most closely associated with improvements in performance. To obtain the EEG activity patterns that most closely corresponded to the integer frequency values of 4, 5, 6, 7, and 8 Hz, we changed the padratio to 8 in further EEG analysis. Comparing the changes of each integer frequency (4 Hz, 5 Hz, 6 Hz, 7 Hz, 8 Hz) activity between block 1 and block 3. Measured the MI between power features (4 Hz, 5 Hz, 6 Hz, 7 Hz, 8 Hz) and n-back performance to investigate which frequency was more sensitive to changes in behavior. Specifically, the frequency with the largest MI magnitude is chosen as the stimulation frequency and was applied to modulate working memory.

### 2.2 Results

#### 2.2.1 Behavioral Analyses

The target-ACC of the n-Back task was analyzed using a mixed-design analysis of variance (ANOVA) employing one between-subject factor of group (HP or LP) and two within-subject factors of back (three-back or four-back) and block (block 1, 2, or 3). As shown in Figure 4, the main effect of block was significant (*F*_2, 38_ = 23.015, *p* =.000, *MSE* =3266.76, *η*^2^ =.55), suggesting that target-ACC increased as the participants practiced more (target-ACC_block3_ > target-ACC_block2_ > target-ACC_block1_, *ps*. <.05). The main effect of back was also significant (*F*_1, 19_ = 25.778,*p* =.000,*MSE* =5831.80, *η*^2^ =.58), suggesting that target-ACC was significantly better on the four-back than the three-back task (*ps*. <.05). The main effect of group was significant (*F*_1, 19_ = 15.003, *p* =.001, *MSE* =5630.21, *η*^2^ =.44), suggesting that target-ACC was significantly better among the HP group than among the LP group (*ps*. <.05). We also observed a significant interaction effect between block and group (*F*_2, 38_ = 6.02, *p* =.005, *MSE* =828.77, *η*^2^ =.24), suggesting that target-ACC increased with practice in the LP group (target-ACCblock3 > target-ACCblock1, target-ACCblock3 > target-ACC_block1_, *ps*. <.05). In the HP group, only block 3 target-ACC was significantly greater than that in block 1. Further comparisons indicated that target-ACC significantly improved as the number of practice trials increased in the LP group (three-back: target-ACC_block3_ > target-ACCblock1,*ps*. <.05; four-back: target-ACCblock3 > target-ACC_block2_ > target-ACC_block1_, *ps*. <.05). However, this effect was not observed in the HP group.

**Figure 3.**
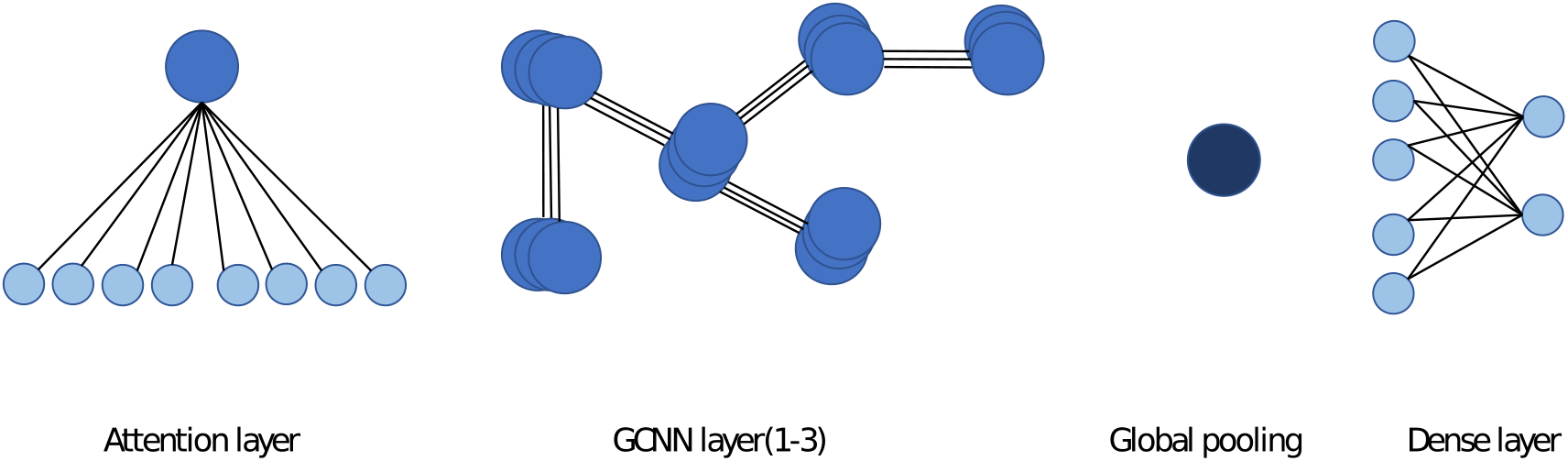
The architecture of the attention based GCNN. The network consists of an attention layer, three GCNN layers, a global pooling layer, and a dense layer. The graph attention mechanism in the first layer learns the dynamic adjacent matrix and the graph features.

**Figure 4.**
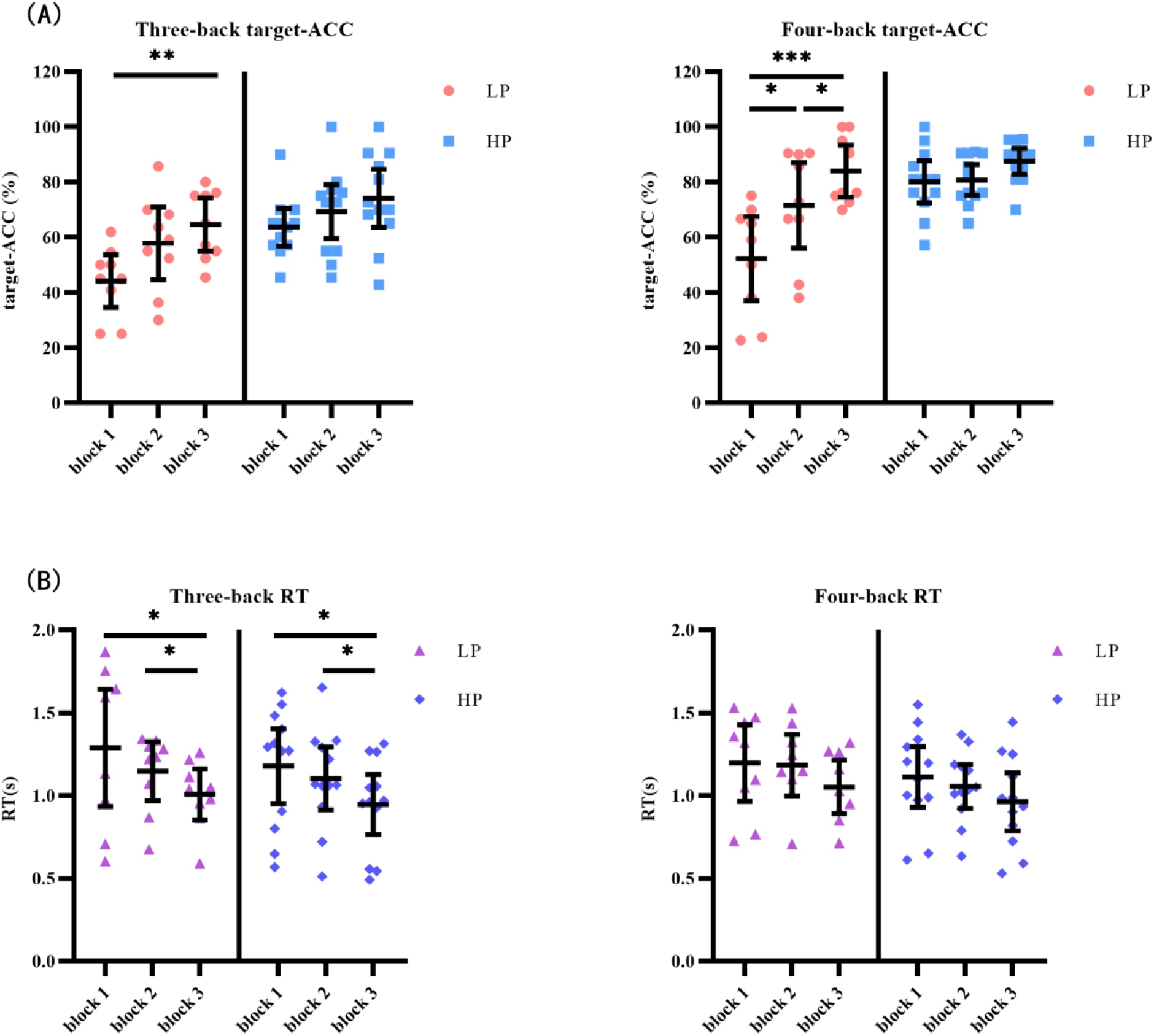
Target-ACC and RT of each group for each back and each block in experiment 1. (A) Scatterplots with individual data points of target-ACC in three-back and four-back tasks. (B) Scatterplots with individual data points of RT in three-back and four-back tasks. Error bars are 95%-confidence intervals around the estimates.

The same mixed-design ANOVA was conducted for the RT of the correct target trials. Only the main effect of block was significant (*F*_1.49, 28.22_ = 13.26, *p* =.000, *MSE* =.43, *η*^2^ =.41, with Greenhouse-Geisser correction), suggesting that the reaction time decreased as the participants practiced more (RT_block1_ > RT_block3_, RT_block2_ > RT_block3_, *ps*. <.05). Further comparisons indicated that RT significantly decreased as the number of practice trials increased in the relatively simple three-back task (For three-back, RT_block1_ > RT_block3_, RT_block2_ > RT_block3_, *ps*. <.05, in both the HP and LP groups), but not in the relatively difficult four-back task.

Our analysis of behavioral outcomes indicates that the target-ACC was affected by naturally capacity of the verbal working memory, and in block 3 the target-ACC was significantly higher than block 1 within the LP group. RT was affected by the difficulty of the task, and the practice effect was only observed in the simpler three-back task.

#### 2.2.2 Initial EEG Analyses

After EEG signal preprocessing, we conducted an initial analysis to explore the brain regions and frequency bands exhibiting changes that corresponded to increases in target-ACC in the LP group. For each frequency band (i.e., theta, alpha, beta) and each block (i.e., block1, block3), the average power between 100 ms and 700 ms was computed and was further averaged among the two n-back tasks. Figure 5 shows the event-related synchronization distribution from block 1 to block 3 (*Power*_*block* 3_ – *Power*_*block* 1_). According to this figure, the power seemed relatively stable in the central and parietal regions, regardless of the group or frequency band. Compared with those in block 1, theta and alpha activity was significantly enhanced in the prefrontal, frontal, and occipital lobes in block 3. Considering that the occipital lobe is more involved in visual processing, while the prefrontal and frontal lobes are more closely related to working memory processing, we focused further analyses on theta and alpha activity in the prefrontal and frontal lobes. After preliminary identification of brain regions and frequencies, the theta and alpha power in Fp1, Fp2, F3, and F4 of the n-back task was analyzed using a mixed-design ANOVA employing one between-subject factor of group (HP or LP) and two within-subject factors of back (three-back or four-back) and block (block 1, 2, or 3). For theta activity, the main effect of block was significant (*F*_2, 34_ = 5.18, *p* =.011, *MSE* =22.99, *η*^2^ =.23) in Fp1, powerblock3 was significantly greater than powerblock1, and powerblock2 was significantly greater than powerblock1. For theta activity, the main effect of block was significant (*F*_2, 34_ = 6.39, *p* =.004, *MSE* =26.12, *η*^2^ =.27) in Fp2, powerblock3 was significantly greater than powerblock1, and powerblock2 was significantly greater than powerblock1. For theta activity, the main effect of block was significant (*F*_2, 34_ = 7.30 *p* =.002, *MSE* =16.79, *η*^2^ =.30) in F3, and power_block2_ was significantly greater than power_block1_. For alpha activity, the main effect of block was significant (*F*_2, 34_ = 3.86 *p* =.031, *MSE* =16.06, *η*^2^ =.19) in Fp2, and power_block3_ was significantly greater than powerblock1. No other main effects were significant (see Table 1). These findings suggested that, when compared with other combinations (i.e., theta in frontal region, alpha in frontal region, alpha in prefrontal region), theta activity in the prefrontal region exhibited trends similar to those observed for changes in behavior (i.e., Compared with block 1, the behavior [target-ACC and RT] and theta activity in block 3 were changed significantly).

**Figure 5.**
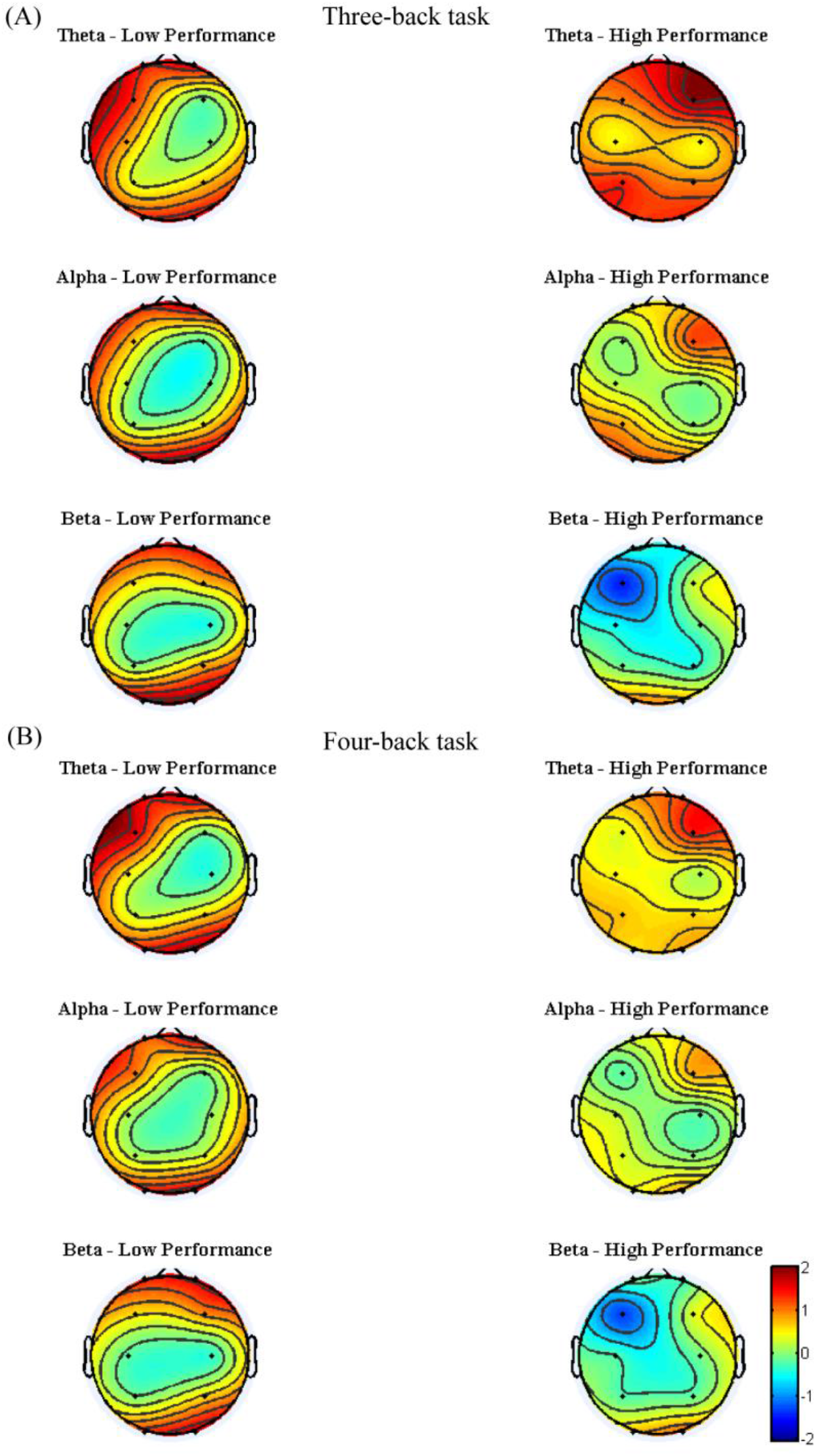
Event-related synchronization from block 1 to block 3 of each group for each band and each back in experiment 1. (A) The power change from block 1 to block 3 in three-back task. (B) The power change from block 1 to block 3 in four-back task. The more tend to red, the more positive changes. The more tend to blue, the more negative changes. The color central region for each group and each frequency band tends to be green, suggesting that the power of central region tend to remain unchanged among practices.

**Table 1.**
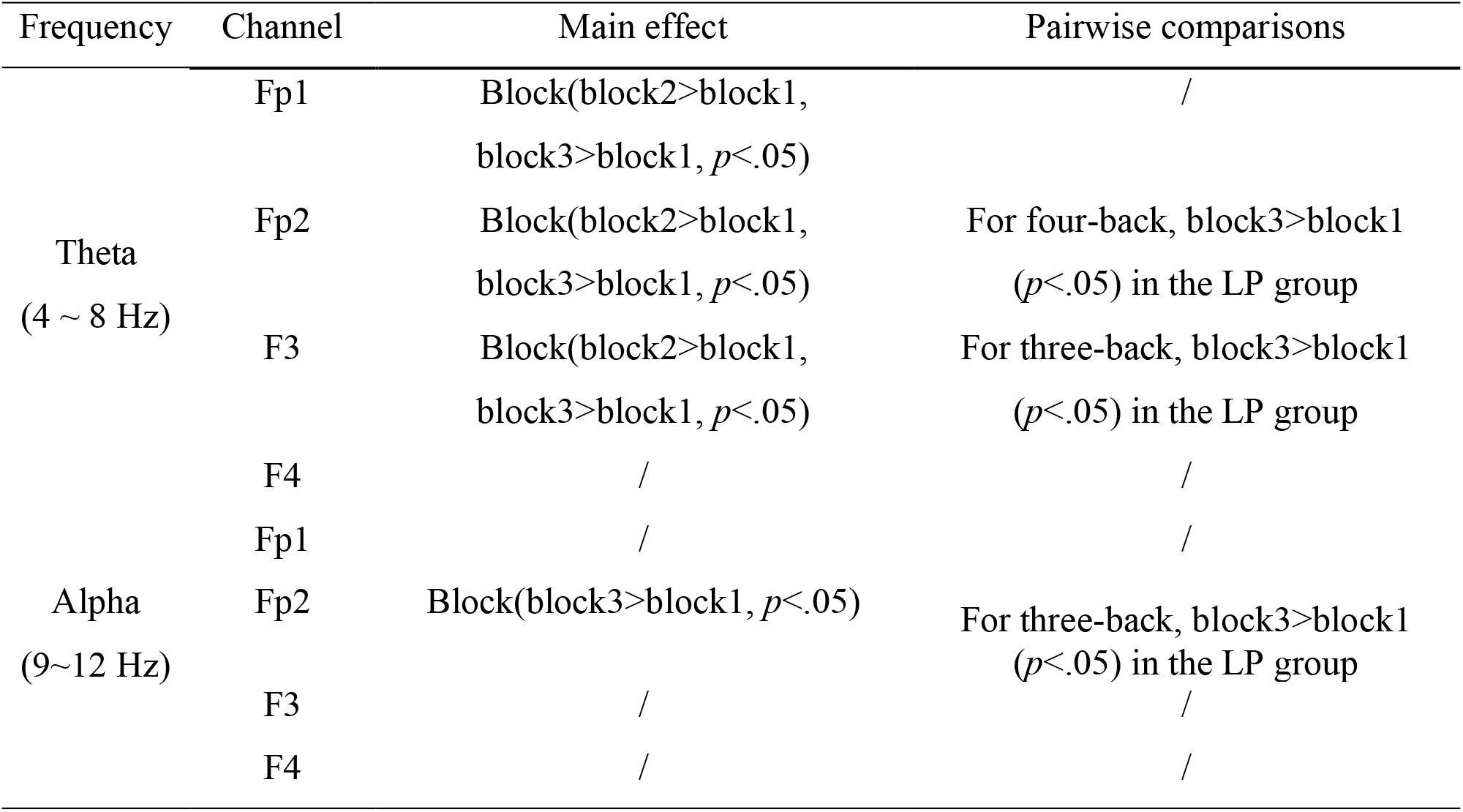
The analysis results of theta and alpha in prefrontal and frontal regions.

Meanwhile, to determine the most discriminative spatio-frequency, we performed quantitative analysis on the 2m (m=2) selected spatial features by measuring MI. We selected the top two largest MI values for each test (see Table 2) and visualized the corresponding EEG topographies (see Figure 6). Table 2 and Figure 6 show that all selected MI values were obtained from the lower band (theta, alpha) activities in frontal and prefrontal region, indicating that lower band activities in frontal and prefrontal region can provide more information for predicting performance on the n-back test. (i.e., more sensitive to n-back performance differences). In particular, among the three tests, the features extracted from the theta band had larger MI values than those extracted from the alpha band, aside from those in the three-back test. Thus, we believe that theta band activity in frontal and prefrontal region may be a better indicator of changes in working memory performance.

**Figure 6.**
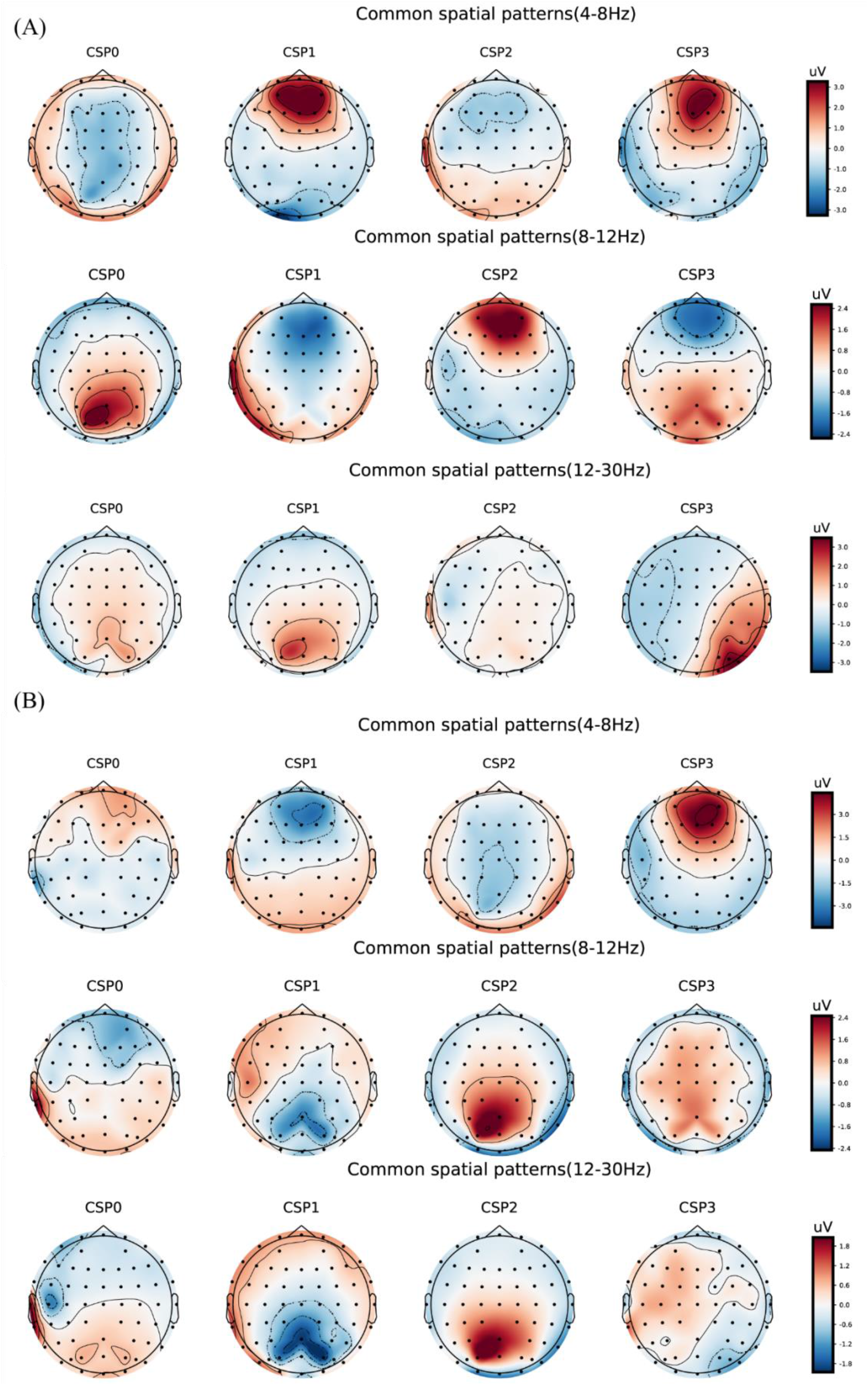
Electroencephalogram topography showing the spatial distribution of the most discriminate features and the associated frequency bands. (A) Electroencephalogram topography of three-back task. From top to bottom, each row displays the most important spatio-frequency modes for the theta, alpha, and beta bands, respectively. (B) Electroencephalogram topography of four-back task.

**Table 2.** The largest MI values in each frequency bands and the corresponding components.

| Test | Component<br>(CSP) | Frequency band | MI |
| --- | --- | --- | --- |
| Three-back | CSP3 | $\theta$ | 0.2677 |
|  | **CSP1** | **$\alpha$** | **0.3264** |
| | CSP0 | $\beta$ | 0.1241 |
| | CSP0 | $\gamma$ | 0.1436 |
| Four-back | **CSP2** | **$\theta$** | **0.2808** |
| | CSP3 | $\alpha$ | 0.2636 |
| | CSP0 | $\beta$ | 0.1395 |
| | CSP0 | $\gamma$ | 0.1921 |

Abbreviations: CSP, common spatial pattern; MI,

#### 2.2.3 Graph Convolutional Neural Network (GCNN)

We use an adapted graph attention mechanism to capture brain network dynamics and find the channel contributing most to performance in the n-back tasks. The proposed model achieved a classification accuracy of 80.4%. We selected the top 15 largest weights from the output optimal adjacent matrix and normalized the chosen weights. The edge between Fp1 and Fp2 had the largest weight at 0.78, suggesting that the functional connection between Fp1 and Fp2 was most important for n-back task performance.

The result of GCNN was similar to the initial EEG analysis, indicating that the brain activity in prefrontal region was associated with the changes in working memory performance.

#### 2.2.4 Further EEG Analysis

In further EEG analysis, we investigated the frequency (4 Hz, 5 Hz, 6 Hz, 7 Hz, and 8 Hz) for which changes in activity were most closely associated with improvements in behavior. The same mixed-design ANOVA was conducted for EEG powers of 4, 5, 6, 7, and 8 Hz in Fp1 and Fp2. Table 3 lists the significant results. We observed that 8 Hz activity in the prefrontal region (especially Fp2) was most closely related to target-ACC. Specifically, for both the three- and four-back tasks, 8 Hz activity was significantly greater in block 3 than in block 1 in the LP group, as was the target-ACC. Prefrontal activity at 6 and 7 Hz appeared to be related to both target-ACC and RT.

**Table 3.** The analysis results of 4 Hz, 5 Hz, 6 Hz, 7 Hz, and 8 Hz in prefrontal.

| Channel | Frequency | Main effect | Pairwise comparisons |
| --- | --- | --- | --- |
| Fp1 | 4 Hz | Block(block2>block1, $p<.05$) | / |
| | 5 Hz | Block(block2>block1, block3>block1, $p<.05$) | / |
| | 6 Hz | Block(block2>block1, block3>block1, $p<.05$) | / |
| | 7 Hz | Block(block2>block1, block3>block1, $p<.05$) | / |
| | 8 Hz | Block(block2>block1, $p<.05$) | / |
| Fp2 | 4 Hz | Block(block3>block1, $p<.05$) | / |
| | 5 Hz | Block(block3>block1, $p<.05$) | / |
| | 6 Hz | Block(block2>block1, block3>block1, $p<.05$) | For three-back, block3>block1 ($p<.05$) in the HP group,<br>For four-back, block3>block1 ($p<.05$) in the LP group |
| | 7 Hz | Block(block2>block1, block3>block1, $p<.05$) | For three-back, block3>block1 ($p<.05$) in the HP group,<br>For four-back, block3>block1 ($p<.05$) in the LP group |
| | 8 Hz | Block(block2>block1, block3>block1, $p<.05$) | For three-back, block3>block1 ($p<.05$) in the LP group,<br>For four-back, block3>block1 ($p<.05$) in the LP group |

Meanwhile, we measured the MI between power features (4 Hz, 5 Hz, 6 Hz, 7 Hz, and 8 Hz) of the two selected regions (Fp1 and Fp2) and n-back performance (see Table 4). We observed that the 8 Hz power of both regions had larger MI values than other frequencies. Since the magnitude of MI is an indicator of shared information between variables, we inferred that dependency was greatest between 8 Hz power and n-back task performance when compared with that for the other four frequency– performance pairs.

**Table 4.** The mutual information between 4 Hz, 5 Hz, 6 Hz, 7 Hz, and 8 Hz power and working memory performance in the two selected regions.

| Channel<br>Frequency (Hz) | Fp1 | Fp2 |
| --- | --- | --- |
| 4 | 0 | 0.002 |
| 5 | 0.006 | 0 |
| 6 | 0 | 0.003 |
| 7 | 0.002 | 0.003 |
| 8 | 0.009 | 0.012 |

Considering the specificity of the stimulus, these findings indicated that applying 8-Hz stimulation to the prefrontal lobe may be effective for improving verbal working memory performance.

#### 2.2.5 Summary

EEG analysis indicated that 8 Hz activity in the prefrontal lobe was associated with the correct response rate in the verbal working memory task, while 6 and 7 Hz activity appeared to be associated with both the correct response rate and response time. Dependency was greatest between 8 Hz power in the prefrontal cortex and n-back task performance when compared with that for the other four frequency–performance pairs. In addition, machine learning results suggested that the functional connection between Fp1 and Fp2 was most important for performance in the n-back tasks. These EEG and machine learning results were used to design Experiment 2, in which the prefrontal lobe was selected as the target for stimulation at a frequency of 8 Hz. In Experiment 2, we compared the modulatory effects of 8 Hz (selected stimulation), 40 Hz (control) and sham stimulation on verbal working memory.

## 3 Experiment 2

### 3.1 Materials and Methods

#### 3.1.1 Participants

In Experiment 2, we recruited 67 young healthy volunteers, but only 48 were included in the behavioral data analysis (20-30 years old). The exclusion criteria were as follows: 1) participants who did not follow the instructions, 2) participants who had outstanding performance in pre-stimulation (target-ACC of >90% in the pre-stimulation tasks), and 3) extreme values (target-ACC or RT exceeding two standard deviations from the mean). Among the 48 included participants, 12 were excluded from the EEG analysis because of poor signal quality, and 36 participants (12 females; mean age 23.67±1.97 years) were included in the analyses.

All participants had normal or corrected-to-normal vision and were right-handed. A preliminary questionnaire screening with each subject ensured that all inclusion criteria for transcranial electric stimulation applications were met (i.e., no history of neuropsychiatric disorders [e.g., epilepsy], no brain injuries, no pregnancy, no intake of neuroleptic or hypnotic medications, and no metallic or electrical implants in the body).

This study was approved by the Ethics Committee of the Shenzhen Institute of Advanced Technology, and all experimental procedures conformed to the principles of the Helsinki Declaration regarding human experimentation. All participants provided oral consent, signed informed consent documents, and received 200RMB for their participation.

#### 3.1.2 Experimental Design and Schedule

Experiment 2 was conducted using a single-blinded sham-controlled design. Participants were randomly divided into a selected group (n = 14), sham group (n = 18), and control group (n = 16). They completed three sessions (pre-stimulation, stimulation, and post-stimulation). Each session included one n-back task (blocks 1, 2, or 3). In the pre-stimulation session, resting-state EEG data were collected for 5 min before the block 1 n-back task. The participants then underwent tACS while performing the block 2 n-back tasks in the stimulation session. The post-stimulation session was the same as that in the pre-stimulation session. Task-state EEG data were recorded for block 1 (the pre-stimulation session) and block 3 (the post-stimulation session). Finally, subjects completed an electrical stimulation sensitivity questionnaire to report their experienced regarding phosphenes, dizziness, tingling, and itching.

#### 3.1.3 N-Back Tasks

Compared with the n-Back tasks of Experiment 1, those in Experiment 2 included an additional five-back task to further investigate the effect of tACS on performance on a more difficult working memory task. In addition, there are nine sequences in total, and each back included three sequences. Each sequence contained 33+n trials, including 11 target trials and 22 non-target trials.

#### 3.1.4 EEG Recordings and Data Preprocessing

EEG data recording, processing, and time-frequency analyses were the same as those in Experiment 1.

#### 3.1.5 Transcranial Alternating Current Stimulation

tACS was delivered via a pair of 4.5 × 5.5 cm^2^ gel electrodes connected to a battery-driven stimulator. The gel electrode impedances were <500 Ω. One of the electrodes was placed over FP1-AP7 and the other was placed over FP2-AF8. The stimulation intensity was 2.0 mA (peak-to-peak current) and was applied for 20 min during the stimulation session in Experiment 2. The selected group received 8 Hz tACS (8 Hz group) and the control group received 40 Hz tACS (40 Hz group). The sham group was also equipped with tACS electrodes but did not receive stimulation.

### 3.2 Results

#### 3.2.1 Behavioral Analyses

The target-ACC of the n-Back task was analyzed using a mixed-design ANOVA employing one between-subject factor of group (8 Hz, 40 Hz, or sham) and two within-subject factors of back (three-back, four-back, or five-back) and block (block 1 or 2). As shown in Figure 7A, the main effect of block was significant (*F*_1, 39_ = 62.56, *p* =.000, *MSE* =6059.908, *η*^2^ =.62), suggesting that target-ACC was greater in block 3 than in block 1. The main effect of back was also significant (*F*_1.56_, 60.87 = 42.90, *p* =.000, *MSE* =611.80, *η*^2^ =.52, with Greenhouse-Geisser correction), suggesting that target-ACC decreased significantly as the difficulty of the task increased (target-ACC_three-back_ > target-ACCfour-back >target-ACC_five-back_, *ps*. <.001). We also observed a significant interaction effect between block and group (*F*_2, 39_ = 7.11, *p* =.002, *MSE* =689.06, *η*^2^ =.27), suggesting that target-ACC was significantly greater for block 3 than for block 1 at 8 Hz, 40 Hz, and in the sham condition (*ps*. <.05). The interaction effect between back and block (*F*_2,78_ = 3.38, *p* =.039, *MSE*=178.69, *η*^2^ =.08) was also significant, suggesting that target-ACC was significantly greater in block 3 than in block 1 for three-back, four-back, and five-back tasks (*ps*. <.05). Further comparisons indicated that block 3 target-ACC in the 8 Hz group was significantly higher than that in block 1 for the three-back, four-back, and five-back tasks (*ps*. <.05). Furthermore, in the sham group, target-ACC was significantly higher in block 3 than in block 1 for the three-back and four-back (*ps*. <.05). However, in the 40 Hz group, target-ACC was significantly greater in block 3 than in block 1 for the three-back task only.

**Figure 7.**
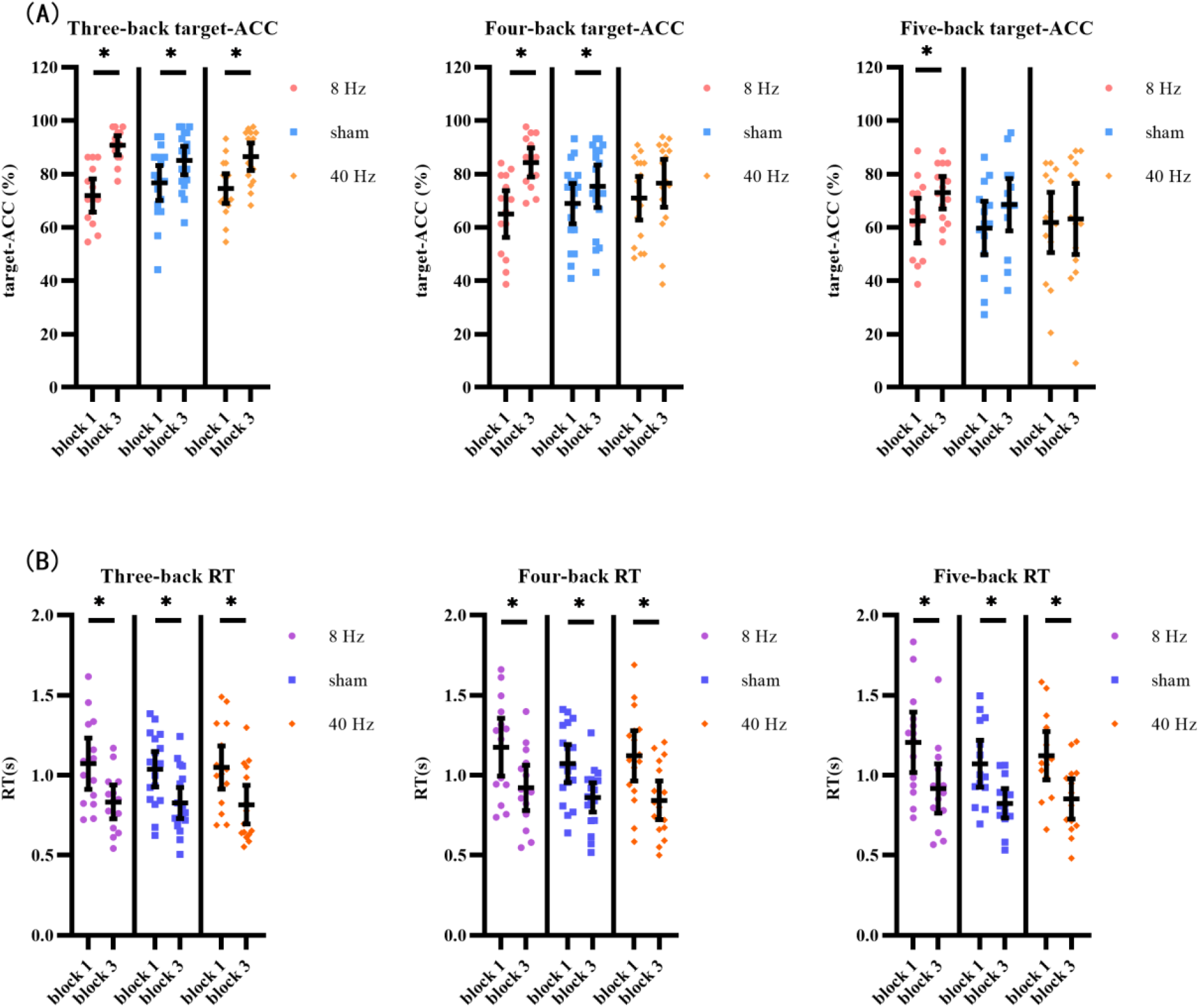
Target-ACC and RT of each group for each back and each block in experiment 2. (A) Scatterplots with individual data points of target-ACC in three-back, four-back, and five-back tasks. (B) Scatterplots with individual data points of RT in three-back, four-back, and five-back tasks. Error bars are 95%-confidence intervals around the estimates.

The same mixed-design ANOVA was conducted to examine RT for correct target trials. As shown in Figure 7B, the main effect of block was significant (*F*_1, 39_ = 87.58, *p* =.000, *MSE* =4.18, *η*^2^ =.69), suggesting that block 3 RTs were shorter than those in block 1. The main effect of back was also significant (*F*_1.59, 62.18_ = 25.12, *p* =.000, *MSE* =.16, *η*^2^ =.39, with Greenhouse-Geisser correction), suggesting that RT increased significantly as the difficulty of the task increased (RTfour-back > RTthree-back, RTfive-back > RTthree-back, *ps*. <.001). Further comparisons indicated that RT was significantly shorter in block 3 than in block 1 for all three groups (8 Hz, 40 Hz and sham) and in all three task conditions (three-back, four-back, and five-back) (*ps*. <.05).

As shown above, the strong practice effect resulted in better performance in block 3 than in block 1. Therefore, we used the improvements in target-ACC (i.e., target-ACC_block3_ – target-ACC_block1_) and RT (i.e., RT_block3_ – RT_block1_) as behavioral indices to compare which stimulation setting induced the greatest improvements in verbal working memory. Improvements in target-ACC in each n-back task were analyzed using a mixed-design ANOVA employing one between-subject factor of group (8 Hz, 40 Hz, or sham) and one within-subject factor of back (three-back, four-back, or five-back). As shown in Figure 8A, the main effect of back was significant (*F*_2, 78_ = 3.38, *p* =.039, *MSE* =357.38, *η*^2^ =.08), indicating a smaller degree of improvement in target-ACC in the five-back task than in the three-back task. The main effect of group was significant (*F*_2, 39_ = 7.11, *p* =.002, *MSE* =1378.12, *η*^2^ =.27), indicating that the target-ACC improvement was significantly greater in the 8 Hz group than in the 40 Hz and sham groups (*ps*. <.05). Further comparisons revealed that the target-ACC improvement of 8 Hz group was significantly greater than that of the 40-Hz group and sham group (*ps*. <.05) in the three-back and four-back tasks. In the five-back task, the improvement in target-ACC was significantly greater in the 8 Hz group than in the 40 Hz group (*ps*. <.05). The same analysis was conducted to examine improvements in RT. However, no significant effects were observed in the RT analysis.

**Figure 8.**
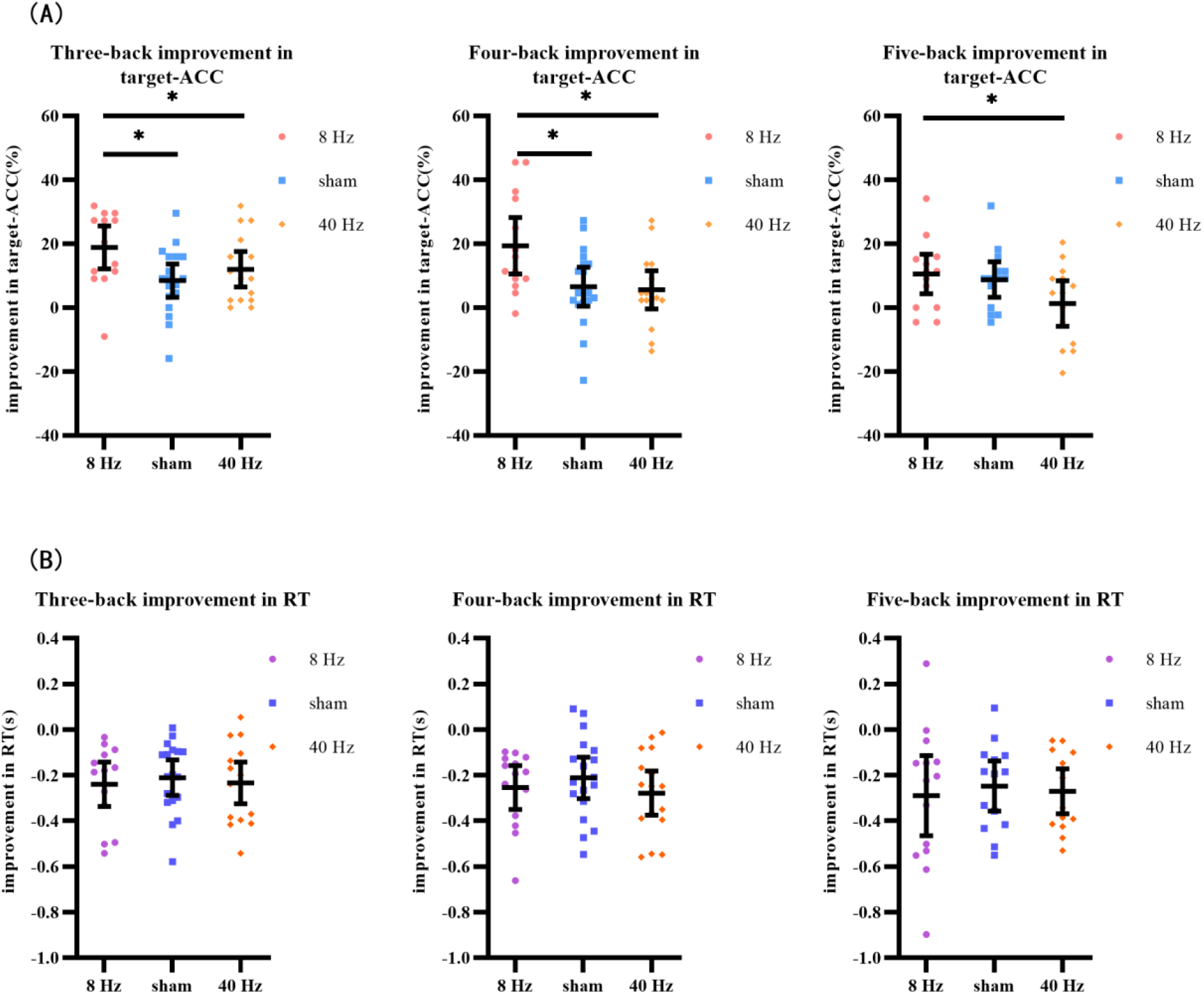
Improvement in target-ACC and RT of each group for each back in experiment 2. (A) Scatterplots with individual data points of improvement in target-ACC for three-back, four-back, and five-back tasks. (B) Scatterplots with individual data points of improvement in RT for three-back, four-back, and five-back tasks. Error bars are 95%-confidence intervals around the estimates.

We further aimed to explore the effects of the three stimulation conditions on verbal working memory in the HP and LP groups, which were determined based on performance in block 1. Scores for the three-back, four-back, and five-back tasks were summed, and participants who scored lower than the median were assigned to the LP group, while those who scored higher than the median were assigned to the HP group. Eventually, the volunteers were divided into six groups: an LP group receiving 8-Hz stimulation (LP-8 Hz) (n = 6), an HP group receiving 8-Hz stimulation (HP-8 Hz) (n = 8), an LP group receiving sham stimulation (LP-sham) (n = 10), an HP group receiving sham stimulation (HP-sham) (n = 8), an LP group receiving 40-Hz stimulation (LP-40 Hz) (n = 8), and an HP group receiving 40-Hz stimulation (HP-40 Hz) (n = 8). The target-ACC of the n-Back task was analyzed using a mixed-design ANOVA employing one between-subject factor of group (LP-8 Hz, HP-8 Hz, LP-sham, HP-sham, LP-40 Hz, and HP-40 Hz) and two within-subject factors of back (three-back, four-back, or five-back) and block (block 1 or 3).

As shown in Figure 9A, the main effect of block was significant (*F*_1, 36_ = 69.90, *p* =.000, *MSE* =6183.12, *η*^2^ =.66), suggesting that target-ACC was significantly greater in block 3 than in block 1. The main effect of back was also significant (*F*_1.49, 53.44_ = 51.48, *p* =.000, *MSE* =6607.74, *η*^2^ =.59, with Greenhouse-Geisser correction), suggesting that target-ACC decreased significantly as the difficulty of the task increased (target-ACCthree-back > target-ACCfour-back > target-ACCfive-back, *ps*. <.001). The main effect of group was significant (*F*_5, 36_ = 19.47, *p* =.000, *MSE* =4190.33, *η*^2^ =.73), suggesting a complex difference between groups. We also observed a significant interaction effect between block and group (*F*_5, 36_ = 4.46, *p* =.003, *MSE* =394.32, *η*^2^ =.38), indicating that target-ACC in block 3 was significantly greater than that in block 1 in both the LP-sham and HP-sham groups (*ps*. <.05). The analysis also indicated that block 3 target-ACC was significantly greater than block 1 target-ACC in the LP-8 Hz and HP-8 Hz groups (*ps*. <.001). Further comparisons indicated that target-ACC in block 3 was significantly greater than that in block 1 in the three-back, four-back, and five-back tasks within the LP-8 Hz group (*ps*. <.05). Within the HP-8 Hz group, block 3 target-ACC was greater than block 1 target-ACC for the three- and four-back tasks only (*ps*. <.05). We also observed improvements in target-ACC between block 1 and 3 of the three-back task in the LP-40 Hz, HP-40 Hz, and HP-sham group (*ps*. <.05). Target-ACC was significantly greater in block 3 than in block 1 for the five-back task in the LP-sham group (*ps*. <.05).

**Figure 9.**
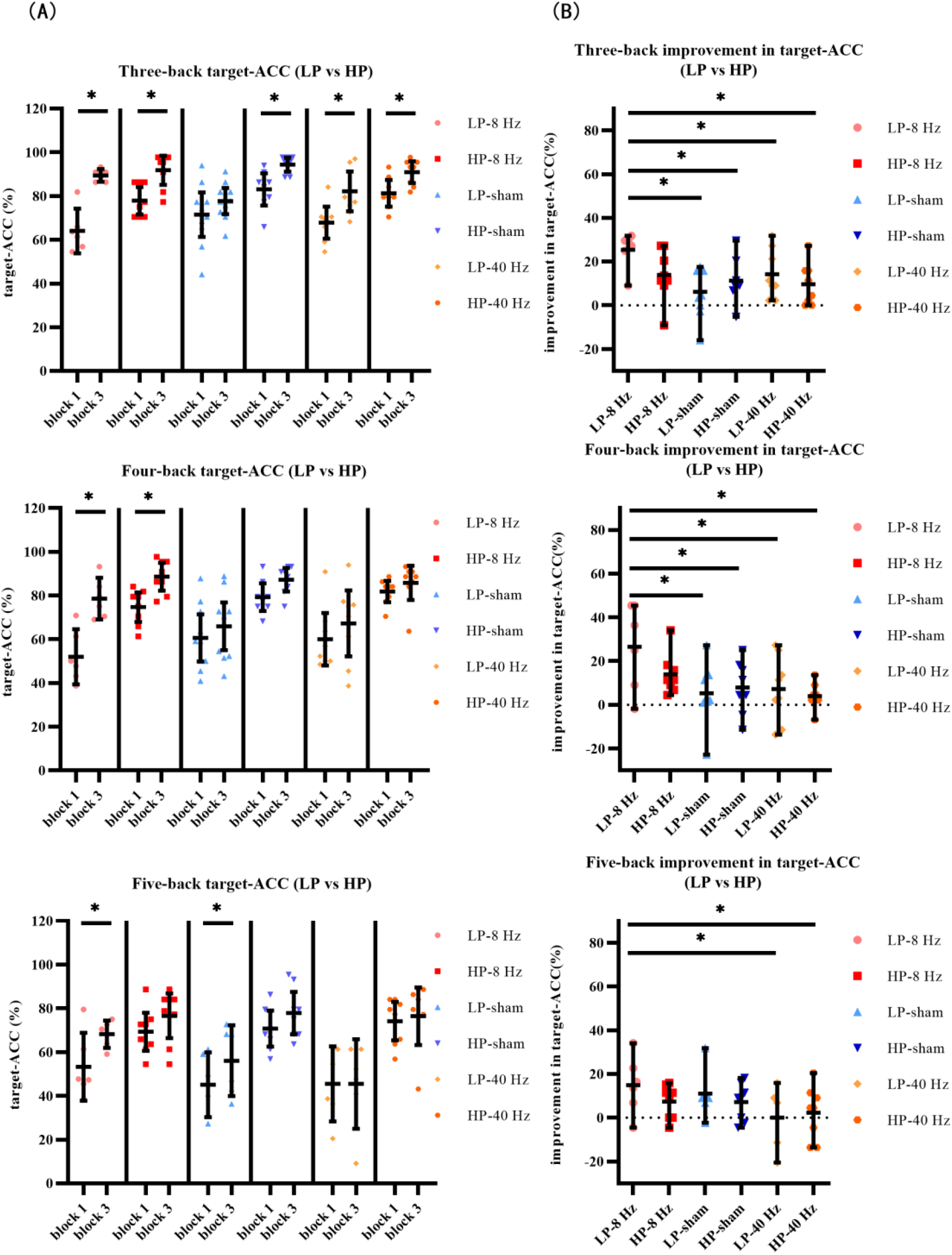
Further analysis result of LP and HP in experiment 2. (A) Scatterplots with individual data points of target-ACC for three-back, four-back, and five-back tasks. (B) Scatterplots with individual data points of improvement in target-ACC for three-back, four-back, and five-back tasks. Error bars are 95%-confidence intervals around the estimates.

To weaken the effect of practice on the results, the target-ACC improvement in the n-back task was analyzed using a mixed-design ANOVA employing one between-subject factor of group (LP-8 Hz, HP-8 Hz, LP-sham, HP-sham, LP-40 Hz, and HP-40 Hz) and one within-subject factor of back (three-back, four-back, or five-back). As shown in Figure 9B, the main effect of group was significant (*F*_5, 36_ = 4.46, *p* =.003, *MSE* =788.64, *η*^2^ =.38), indicating that the target-ACC improvement was significantly greater in the LP-8 Hz group than in the other groups. Further comparisons revealed that the target-ACC improvement was significantly greater in the LP-8 Hz group than in the LP-sham, HP-sham, LP-40 Hz, and HP-40 Hz groups in the three-back and four-back tasks (*ps*. <.05). In the five-back task, the target-ACC improvement of the LP-8 Hz group was significantly greater than that of the LP-40 Hz and HP-40 Hz groups (*ps*. <.05). As our previous analysis revealed no differences in the effects of the three stimulation conditions on RT, we did not analyze RT results here.

#### 3.2.2 EEG Analyses

The theta power in Fp1 and Fp2 during the n-back task was analyzed using a mixed-design ANOVA employing one between-subject factor of group (8 Hz, 40 Hz, or sham) and two within-subject factors of back (three-back, four-back, or five-back) and block (block 1 or 2). In Fp1, the main effect of block was significant (*F*_1, 33_ = 4.23, *p* =.048, *MSE* =95.32, *η*^2^ =.11), and the theta power was significantly greater in block 3 than in block 1. Further comparisons indicated that theta power during the three-back and four-back tasks was significantly greater in block 3 than in block 1 in the 8 Hz group (*ps*. <.05), while that during the five-back task was only marginally significantly greater (*ps*.=.056). No significant effects were observed in the 40 Hz and sham groups. In Fp2, the main effect of block was marginal significant (*ps*. = .056), and the theta power was greater in block 3 than in block 1. Furthermore, no effect was significant in the theta power analysis for Fp2.

In addition, we explored the effects of the three stimulation conditions on EEG activity associated with verbal working memory in the HP and LP groups. The theta power in Fp1 and Fp2 during the n-back task was analyzed using a mixed-design ANOVA employing one between-subject factor of group (LP-8 Hz, HP-8 Hz, LP-sham, HP-sham, LP-40 Hz, and HP-40 Hz) and two within-subject factors of back (three-back, four-back, or five-back) and block (block 1 or 2). No effect was significant in the theta power analysis for either Fp1 or Fp2.

#### 3.2.3 Adverse Effects Ratings

Participants were required to rate their adverse experiences during and after stimulation. The questionnaire used a four-point Likert scale ranging from 1 (none) to 4 (extreme). Overall, tACS was well-tolerated. For 8-Hz and 40-Hz stimulation, participants reported phosphenes (100%), dizziness (40%), tingling (73.33%), and itching (46.67%) during stimulation. These effects were attenuated after the stimulation, and participants reported phosphenes (3.45%), dizziness (24.14%), tingling (0%), and itching (6.9%). According to the one-way ANOVA, most of the ratings of adverse experiences that occurred during stimulation significantly differed between groups, including phosphenes (*F*_2, 45_ = 20.52, *p*<.001), tingling (*F*_2, 45_ =14.15, *p*<.001), and itching (*F*_2, 45_ = 3.71, *p* =.032). For phosphenes and tingling, the ratings of the 8 Hz and 40 Hz groups were significantly greater than those of the sham group (*ps* <.001). For itching, the rating of the 40 Hz group was significantly greater than that of the sham group (*p* =.034). However, the rating of dizziness that occurred during stimulation did not significantly differ between the groups (*F*_2, 45_ = 1.52, *p* =.230). The ratings of adverse experiences that occurred after stimulation did not significantly differ between groups.

## 4 Discussion

### 4.1 EEG activities related to positive behavior changes

In Experiment 1, participants complete three n-back tasks (blocks 1, 2, and 3). One week between block 1 and block 2. Ten minutes between block 2 and block 3. The result showed that the practice effect was not affected by the interval time. Practice effect of target-ACC was mainly affected by participant’s naturally verbal working memory capacity. Low performance subjects showed stronger practice effects than high performance participants. Practice effect of RT was mainly affected by the task difficulty. Subjects showed stronger practice effect in relatively simple three-back task than in relatively difficult four-back task.

In initial EEG analysis, we first locked the EEG characteristic regions and frequency bands by observing the differences of the topographic maps between block 3 and block 1. We found that theta and alpha activation of block 3 was greater than block 1 in prefrontal and frontal regions. Specifically, theta activity in the prefrontal region exhibited trends similar to those observed for changes in behavior. Meanwhile, the result of FBCSP suggested that theta band activity in frontal and prefrontal regions may be a better indicator of changes in working memory performance. In addition, we used an adapted graph attention mechanism to capture the brain network dynamics and to find the channel contributing most to the performance in n-back tasks. The result was similar to EEG analysis, finding that brain activity in prefrontal region was associated with the changes in working memory performance. Thus, we concluded that theta activity in prefrontal region was associated with improvements in verbal working memory performance.

In further EEG analysis, we investigated the frequency (4 Hz, 5 Hz, 6 Hz, 7 Hz, and 8 Hz) for which changes in activity were most closely associated with improvements in behavior. The result indicated that 8 Hz activity in the prefrontal lobe was associated with the correct response rate in the verbal working memory task, while 6 and 7 Hz activity appeared to be associated with both the correct response rate and response time. Considering the specificity of the stimulus, these findings indicated that applying 8-Hz stimulation to the prefrontal lobe may be effective for improving verbal working memory performance.

### 4.2 The modulatory effects of 8 Hz (selected stimulation), 40 Hz (control), and sham stimulation on verbal working memory

In experiment 2, we compared the modulatory effects of 8 Hz (selected stimulation), 40 Hz (control), and sham stimulation on verbal working memory. The strong practice effects showed better performance in block 3 than block 1. Therefore, we used the improvements in target-ACC and RT as behavioral indices to compare which stimulation setting induced the greatest improvements in verbal working memory. The target-ACC improvement of 8 Hz group was significantly greater than that 40 Hz group and sham group in the three-back and four-back tasks. However, no significant effects were observed in RT analysis. Those results confirmed to the inference of experiment 1, 8 Hz activity in prefrontal region was associated with the correct response rate in verbal working memory task.

We further explored the effects of three stimulation conditions on verbal working memory in HP and LP groups, which were determined based on performance in block 1. In a relatively simple three-back task, target-ACC for most of subjects had a significantly higher in block 3 than block 1. In relatively difficult four-back and five-back tasks, only LP-8 Hz group maintained a stable and significant improvement in target-ACC. The improvements in target-ACC of LP-8 Hz was significantly greater than 40 Hz and sham group. The target-ACC of verbal working memory was improved significantly using 8 Hz stimulation than 40 Hz and sham stimulation (Especially for participants with low verbal working memory).

Overall, accordance to with several previous studies (Biel et al., 2021; Kilian et al., 2020; Pahor & Jaušovec, 2018; Vosskuhl et al., 2015), our findings indicated that theta band activity was strongly associated with verbal working memory, and that theta tACS improved verbal working memory performance. Moreover, our study extends these findings, as we investigating the EEG characteristics correspond to the improvements in working memory performance in both HP and LP groups. Our analysis revealed that the changes in 8 Hz activity prefrontal region exhibited trends similar to those for the correct response rate in verbal working memory tasks. These results may indicate that 8 Hz activity in prefrontal region supports response accuracy. In Experiment 2, we applied 8 Hz tACS in prefrontal region, representing the biggest difference between the current investigation and previous studies. Although our stimulus targets and frequencies differed from those used in previous research, the performance of verbal working memory was improved significantly by using 8 Hz stimulation than 40 Hz and sham stimulation (Especially for participants with low verbal working memory). The result suggested that 8 Hz tACS at prefrontal region had an effective intervention on improving verbal working memory.

In EEG analysis, the theta power of prefrontal region during n-back tasks was greater in block 3 than block 1 whith 8 Hz group. No significant effects were observed in 40 Hz and sham groups. The results suggested that after stimulation 8 Hz tACS improves brain oscillations of the theta frequency band. It is worth considering that the degree of change in EEG is relatively subtle compared to the change in behavior.

### 4.3 Conclusions

The results of Experiment 1 showed that prefrontal lobe theta power was particularly sensitive to the amount of practice. Specifically, 8 Hz activity in the prefrontal region was related to improvements in response accuracy among participants with low verbal working memory ability, while activity at 6 and 7 Hz was related to both response accuracy and RT. Meanwhile, machine learning also indicated that frontal lobe theta power (especially for 8 Hz activity) is sensitive to improvements in performance. In Experiment 2, we utilized a frequency of 8 Hz to target the prefrontal region during tACS. The results of Experiment 2 showed that 8 Hz tACS could effectively improve performance on verbal n-back tasks, and the brain oscillations of the theta frequency band increased after stimulation. In addition, when 8 Hz stimulation was delivered, the target-ACC improvement was significantly higher in the LP group than other participants in sham and 40 Hz groups. These results suggest that applying 8 Hz electrical stimulation to the prefrontal region can effectively improve verbal working memory performance (especially in individuals with low ability), while stimulation at 40 Hz and sham stimulation exert no such effects. In conclusion, using EEG features related to positive behavioral changes to select brain regions and stimulation patterns for tACS is an effective intervention for improving working memory.

### 4.4 Significance

The current study indicated that employing 8 Hz tACS in the prefrontal region can improve performance on n-back tasks that assess working memory. Delivery of tACS at 8 Hz may be especially helpful for improving verbal working memory in participants with generally low initial ability. Moreover, few studies to date have focused on stimulation at Fp1 and Fp2.

More importantly, the current study provides new insight into the selection of appropriate parameters for tACS. Researchers can first investigate the neurophysiological features associated with positive behavioral changes in specific cognitive tasks. Then, selecting tACS targets and parameters based on the feature. This method could be particularly helpful when the source of brain oscillations of specific cognitive functions is not clearly understood. For example, most tACS can influence the superficial regions of brain cortex only (Brunyé, 2018). The spatial resolution of EEG is relatively low. If stimulation of superficial brain regions can causally influence cognitive function, the experiment could indicate that the specific superficial brain region is involved in cognitive function. Using the same logic, different combinations of neurophysiological and stimulation approaches can also be employed to study the mechanisms of cognitive functions and aid the development of interventions for various mental disorders.

### 4.5 Limitations

The current study applied various analyses to determine the EEG features associated with improvements in verbal working memory performance. The results of these analyses were not homogeneous. The final selection of the parameters was a balance between the results of these analyses. Therefore, the selection of the parameters was not stable, meaning that selection may depend on the number and types of analyses used. If more analyses are included, the results may be more inconsistent, which may make selection difficult. However, the number of analyses was not a key feature of the current paradigm. It is important to determine the parameters of tACS by analyzing the electrophysiological online signal, regardless of the number of analyses employed. In addition, the target region for stimulation was very large and may have covered at least four channel sites in the EEG cap. This shortcoming was mainly attributed to the tACS design. The more specific the region, the smaller the electrode, and the more pain the participants would experience. This pain could drastically reduce cognitive function because it constitutes a significant distraction.

Finally, the approach utilized in the current study may be inconvenient because it requires at least two separate experiments. It has been suggested that individualized stimulation may be better. For example, researchers could analyze the neurophysiological data for each participant immediately after the first test of cognitive function and immediately apply the stimulation in the same experiment. In this scenario, the difference between correct and incorrect trials could be revealed by rapid analyses or machine learning methods. However, these issues are much more complicated in practice. For example, correct trials do not fully reflect true judgment; participants may press a button based on guesswork. Although researchers could subtract the false alarm rate from the target-ACC to evaluate function, the number of correct trials wherein participants respond by guessing would be unknown. Including these trials in the analyses will greatly reduce the reliability of the analyses that aim to differentiate correct and incorrect trials because different participants might have different tendencies to guess. In addition, the number of correct and incorrect trials is difficult to control, and they would directly influence the results of the analyses.

### 4.6 Further Study

Further studies could employ the same procedures in a cohort of older adults to investigate whether this method is effective in improving the working memory of the older adults and those with cognitive decline.

In addition, more cognitive tasks that are used to assess working memory could be included in future studies, employing a similar design to Experiments 1 and 2. By doing this, the differences and the common brain activities of working memory among various tasks could be revealed, which may help to explain the inconsistent results of previous studies.

Future studies should employ AI training to improve cognitive function. Although the differentiation between correct and incorrect trials may be difficult, differentiation of HP and LP groups using AI is feasible. The current study already trained AI to differentiate the two groups. In further studies, this AI could analyze all trials in the first session and classify the case as HP or LP. In the second session, half of the cases in each group would receive the corresponding stimulation. Comparison of the stimulated and non-stimulated cases in the LP group may be more convincing because the two sessions would include the same participants and comparison within a group might make the difference greater.

## 6 Author Contributions

PW designed the study. MG and LZ contributed to the literature search, data collection, data analysis, and interpretation of the results. YL provided hardware support for the experiment and participated in data collection. RW is responsible for machine learning data analysis. All authors contributed to the writing of this paper.

## 8 Acknowledgements

This work was supported in part by the National Key R&D Program of China (2018YFA0701403 to P.W.), the Youth Innovation Promotion Association of the Chinese Academy of Sciences (2017413 to P.W.).

## 9 Data availability statement

The authors declare that the data supporting the findings of this study are available within the article or from the authors upon request.

## References

Ali, M. M., Sellers, K. K., and Fröhlich, F. (2013). Transcranial alternating current stimulation modulates Large-Scale cortical network activity by network resonance. J Neurosci. 33, 11262–11275. https://doi.org/10.1523/JNEUROSCI.5867-12.2013

Ang, K. K., Chin, Z. Y., Zhang, H., and Guan, C. (2008). Filter bank common spatial pattern (FBCSP) in brain-computer interface. In 2008 IEEE international joint conference on neural networks (IEEE world congress on computational intelligence) (pp. 2390–2397). IEEE.

Babiloni, C., Lizio, R., Marzano, N., Capotosto, P., Soricelli, A., Triggiani, A. I., Cordone, S., Gesualdo, L., and Del Percio, C. (2016). Brain neural synchronization and functional coupling in Alzheimer’s disease as revealed by resting state EEG rhythms. Int J Psychophysiol. 103, 88–102. https://doi.org/10.1016/j.ijpsycho.2015.02.008

Battiti, R. (1994). Using mutual information for selecting features in supervised neural net learning. IEEE Trans Neural Netw. 5, 537–550. doi: 10.1109/72.298224.

Benussi, A., Cantoni, V., Cotelli, M. S., Cotelli, M., Brattini, C., Datta, A., Thomas, C., Santarnecchi, E., Pascual-Leone, A., and Borroni, B. (2021). Exposure to gamma tACS in Alzheimer’s disease: A randomized, double-blind, sham-controlled, crossover, pilot study. Brain Stimul. 14, 531–540. https://doi.org/10.1016/j.brs.2021.03.007

Blankertz, B., Dornhege, G., Krauledat, M., Müller, K. R., & Curio, G. (2007). The non-invasive Berlin Brain–Computer Interface: fast acquisition of effective performance in untrained subjects. NeuroImage. 37, 539–550. doi: 10.1016/j.neuroimage.2007.01.051.

Brainard DH (1997). The Psychophysics Toolbox. Spat Vis. 10:433–436.

Brunyé, T. T. (2018). Modulating Spatial Processes and Navigation via Transcranial Electrical Stimulation: A Mini Review. Front Hum Neurosci. 11, 649. doi: 10.3389/fnhum.2017.00649.

Biel, A. L., Sterner, E., Röll, L., & Sauseng, P. (2021). Modulating verbal working memory with fronto-parietal transcranial electric stimulation at theta frequency: Does it work? European Journal of Neuroscience, 1–21. https://doi.org/10.1111/ejn.15563

Chung, Fan RK, and Fan Chung Graham. Spectral graph theory. No. 92. American Mathematical Soc., 1997

Cover, T. M. (1999). Elements of information theory. John Wiley & Sons.

Grover, S., Nguyen, J. A., Viswanathan, V., & Reinhart, R. M. (2021). High-frequency neuromodulation improves obsessive–compulsive behavior. Nat Med. 27, 232–238. doi: 10.1038/s41591-020-01173-w.

Hoy, K. E., Bailey, N., Arnold, S., Windsor, K., John, J., Daskalakis, Z. J., Fitzgerald, P. B. (2015). The effect of γ-tACS on working memory performance in healthy controls. Brain Cogn. 101, 51–56. doi: 10.1016/j.bandc.2015.11.002.

Kirchner, W. K. (1958). Age Differences in Short-Term Retention of Rapidly Changing Information. J Exp Psychol. 55, 352–358. doi: 10.1037/h0043688.

Klink, K., Peter, J., Wyss, P., and Klöppel, S. (2020). Transcranial Electric Current Stimulation During Associative Memory Encoding: Comparing tACS and tDCS Effects in Healthy Aging. Front Aging Neurosci. 12, 66. https://doi.org/10.3389/fnagi.2020.00066

Koenig, T., Prichep, L., Dierks, T., Hubl, D., Wahlund, L. O., John, E. R., and Jelic, V. (2005). Decreased EEG synchronization in Alzheimer’s disease and mild cognitive impairment. Neurobiol Aging. 26, 165–171. doi: 10.1016/j.neurobiolaging.2004.03.008.

Krebs, C., Peter, J., Wyss, P., Brem, A. K., and Klöppel, S. (2021). Transcranial electrical stimulation improves cognitive training effects in healthy elderly adults with low cognitive performance. Clin Neurophysiol. 132, 1254–1263. https://doi.org/10.1016/j.clinph.2021.01.034

Li, S. C., Lindenberger, U., and Sikström, S. (2001). Aging cognition: from neuromodulation to representation. Tren Cogn Sci. 5, 479–486. doi: 10.1016/s1364-6613(00)01769-1.

Misselhorn, J., Göschl, F., Higgen, F. L., Hummel, F. C., Gerloff, C., and Engel, A. K. (2020). Sensory capability and information integration independently explain the cognitive status of healthy older adults. Sci Rep. 10. https://doi.org/10.1038/s41598-020-80069-8

Pahor, A., and Jaušovec, N. (2018). The effects of theta and gamma tacs on working memory and electrophysiology. Front Hum Neurosci. 11:651. https://doi.org/10.3389/fnhum.2017.00651

Pelli DG (1997) The Video Toolbox software for visual psychophysics: trians-forming numbers into movies. Spat Vis. 10:437–442.

Pfurtscheller, G., and Neuper, C. (2001). Motor imagery and direct brain-computer communication. Proceedings of the IEEE. 89, 1123–1134.

Reinhart, R. M. G., and Nguyen, J. A. (2019). Working memory revived in older adults by synchronizing rhythmic brain circuits. Nat Neurosci. 22, 820–827. https://doi.org/10.1038/s41593-019-0371-x

Riddle, J., McFerren, A., and Frohlich, F. (2021). Causal role of cross-frequency coupling in distinct components of cognitive control. Prog Neurobiol. 202, 102033. doi: 10.1016/j.pneurobio.2021.102033.

Rombouts, S. A., Barkhof, F., Goekoop, R., Stam, C. J., and Scheltens, P. (2005). Altered resting state networks in mild cognitive impairment and mild Alzheimer’s disease: an fMRI study. Hum Brain Mapp. 26, 231–239. https://doi.org/10.1002/hbm.20160

Timme, Nicholas M., and Christopher Lapish. “A tutorial for information theory in neuroscience.” eneuro 5.3 (2018).

Vandergheynst P. Convolutional neural networks on graphs with fast localized spectral filtering

Veličković P, Cucurull G, Casanova A, et al. Graph attention networks [J]. arXiv preprint arXiv:1710.10903, 2017

Vosskuhl, J., Huster, R. J., and Herrmann, C. S. (2015). Increase in short-term memory capacity induced by down-regulating individual theta frequency via transcranial alternating current stimulation. Front Human Neurosci. 9, 257. doi: 10.3389/fnhum.2015.00257.

Wang, P., Göschl, F., Friese, U., Engel, A. K. (2019) Long-range functional coupling predicts performance: Oscillatory EEG networks in multisensory processing. Neuroimage 196, 114–125. doi: 10.1016/j.neuroimage.2019.04.001.

Zaehle, T., Rach, S., and Herrmann, C. S. (2010). Transcranial alternating current stimulation enhances individual alpha activity in human EEG. PloS one, 5, e13766. doi: 10.1371/journal.pone.0013766.

